# Expression of *ppsD*, a gene involved in synthesis of *Mycobacterium tuberculosis* virulence factor PDIM, reflects treatment response in pulmonary tuberculosis patients

**DOI:** 10.1101/576470

**Authors:** Kalpana Sriraman, Rupali Kekane, Daksha Shah, Dhananjaya Saranath, Nerges Mistry

**Author notes:** **Corresponding author.**; / 8601.

## Abstract

Drug resistant tuberculosis (TB) cases are primarily driven by transmission, however, treatment failure and acquisition of drug resistance are still significant issues in drug sensitive TB cases. Study of gene expression in *Mycobacterium tuberculosis* (Mtb) isolated from poor outcome patients may offer clues towards prediction of treatment response. In the current study, expression of five non-drug target genes (*ppsD, embC, Rv1457c, Rv1687c* and *recB)* previously identified to be associated with drug resistance was studied in clinical isolates from patients with different treatment outcomes to examine its correlation to treatment response and acquisition of drug resistance in Mtb. Our results show that expression of *ppsD*, a gene involved in synthesis of cell wall lipid PDIM, was significantly increased in patients who developed drug resistance during treatment and patients who were drug resistant at diagnosis. On the other hand in longitudinal isolates collected during treatment, *ppsD* expression decreased consistently in patients who responded to treatment and became culture negative, while it increased in patients who did not respond to treatment as indicated by their culture positive status towards the end of treatment. These results demonstrate that *ppsD* expression reflects treatment response in TB patients and hence can be potentially used as a marker for predicting treatment response. Additional longitudinal studies with a larger cohort of patients are required to establish application of *ppsD* expression as a marker of treatment response.

## Introduction

India has the highest burden of tuberculosis (TB) with 2.8 million new cases reported in 2016[1]. According to India Annual TB report treatment outcome data for microbiologically confirmed cases in 2015[2], about 7% of new TB patients and 16% of retreatment TB patients either failed treatment or developed resistance or died despite treatment. In some states, the percentages are higher than the national average[2]. These patients represent the poor-responders of treatment, essentially the bacteria they harbor do not respond to treatment because of tolerance or resistance to anti-TB drugs. Although the percentage appears to be small, considering the number of estimated cases, it is substantially significant in terms of absolute numbers to be ignored. Currently, methods to predict a priori patients that will respond to treatment or those who will develop drug resistance and respond poorly to treatment are lacking. Studying the bacterial physiology in poor outcome patients may offer clues towards prediction of treatment response.

*Mycobacterium tuberculosis* (Mtb) has developed several mechanisms to combat drug pressure. The most stable mechanism is development of mutations in drug-specific target genes that make the drug ineffective[3]. Other alternate mechanisms like low cell wall permeability, modifying enzymes and efflux offers generalized protection against drug pressure[4,5]. In addition, several studies have shown that these alternative mechanisms sequentially assist in the development of a stable drug resistance mechanism[4]. Studies targeting these mechanisms to improve the effectivity of existing treatments are available. Efflux inhibitors are shown to reduce the minimum inhibitory concentration (MIC) and increase susceptibility to drugs, thus showing promise as adjunct therapeuty[6]. Recent literature shows these mechanisms act in conjunction with drug resistant mutations to determine the ability of bacteria to respond to drugs [7]. Machado et.al.[7] showed that clinical strains with same mutation conferring antibiotic resistance had different MICs and this differences can be reduced by using efflux inhibitors. The contribution of these alternative drug resistance mechanisms to bacterial survival under drug pressure, therefore, cannot be overlooked making it pertinent to understand the role of these mechanisms.

Studying gene expression changes may indicate how these mechanisms relate to the development of drug resistance or response to treatment. Studies on global expression changes in longitudinal clinical isolates from patients who develop drug resistance during treatment and in drug resistant isolates from patients who were resistant at diagnosis have indicated potential candidate genes to predict drug resistance in clinical strains[8,9]. The studies have also shown that deregulation of DNA repair mechanisms may have an influence on mutational abilities of the bacteria and hence enhance the bacterial ability to acquire specific drug target mutations. In an earlier publication, we systematically analyzed these changes using *in vitro* generated mono and multi-drug resistant strains and identified 5 critical genes that were not previously known to be associated with the development of drug resistance[10]. The 5 genes were *ppsD* and *embC* belonging to cell wall processes that determine cell permeability, *recB* belonging to DNA repair, *Rv1457c* and *Rv1687c* that are putative efflux proteins. In this study, we studied the expression of these five genes in clinical isolates from patients with different treatment outcomes to investigate its correlation to treatment response and acquisition of drug resistance in Mtb.

## Materials and Methods

### Study Population and Sample collection

This study was approved by Institutional Ethics Committee (FMR/IEC/TB/02/2015), Central TB Division-New Delhi and Brihanmumbai Municipal Corporation-Mumbai. A total of 168 smear positive sputum samples (baseline) of new pulmonary TB patients were collected from Revised National TB Control Programme (RNTCP) designated microscopy centers (DMC) in two municipal wards of Mumbai between Jan 2016 andJan 2017. The study researchers did not have any direct contact with patients. All samples were subjected to GeneXpert testing. We obtained 132 drug sensitive (DS), 23 rifampicin resistant TB patients and 13 Mtb negative by GeneXpert. The study design involved follow-up (FU) of all DS-TB patients till the end of treatment with sputum sample collected at 2 months (end of intensive phase-FU-1), 4 months (FU-2) and 6 months of treatment (end of treatment-FU-3). Of the 132 DS patients, only 63 patients could be followed up till the end of treatment and at least two follow-up samples per patient were collected. Details of sample collection are described in Figure 1. Sixty nine patients were lost to follow-up with reasons being-defaulter (18), expired (11), initial defaulter/shifted to private care(15), referred out to a different health post(12), migrated (9) or others (4). Of the 69 lost to FU patients who were alive, treatment outcome information at 6 months of treatment could be traced for 4 patients. By design, patients detected as DR at baseline were not followed up, however, we received random follow-up samples (4^th^, 6^th^ and 18^th^ month) of one patient from DMC, which were used for gene expression analysis.

**Fig 1:**
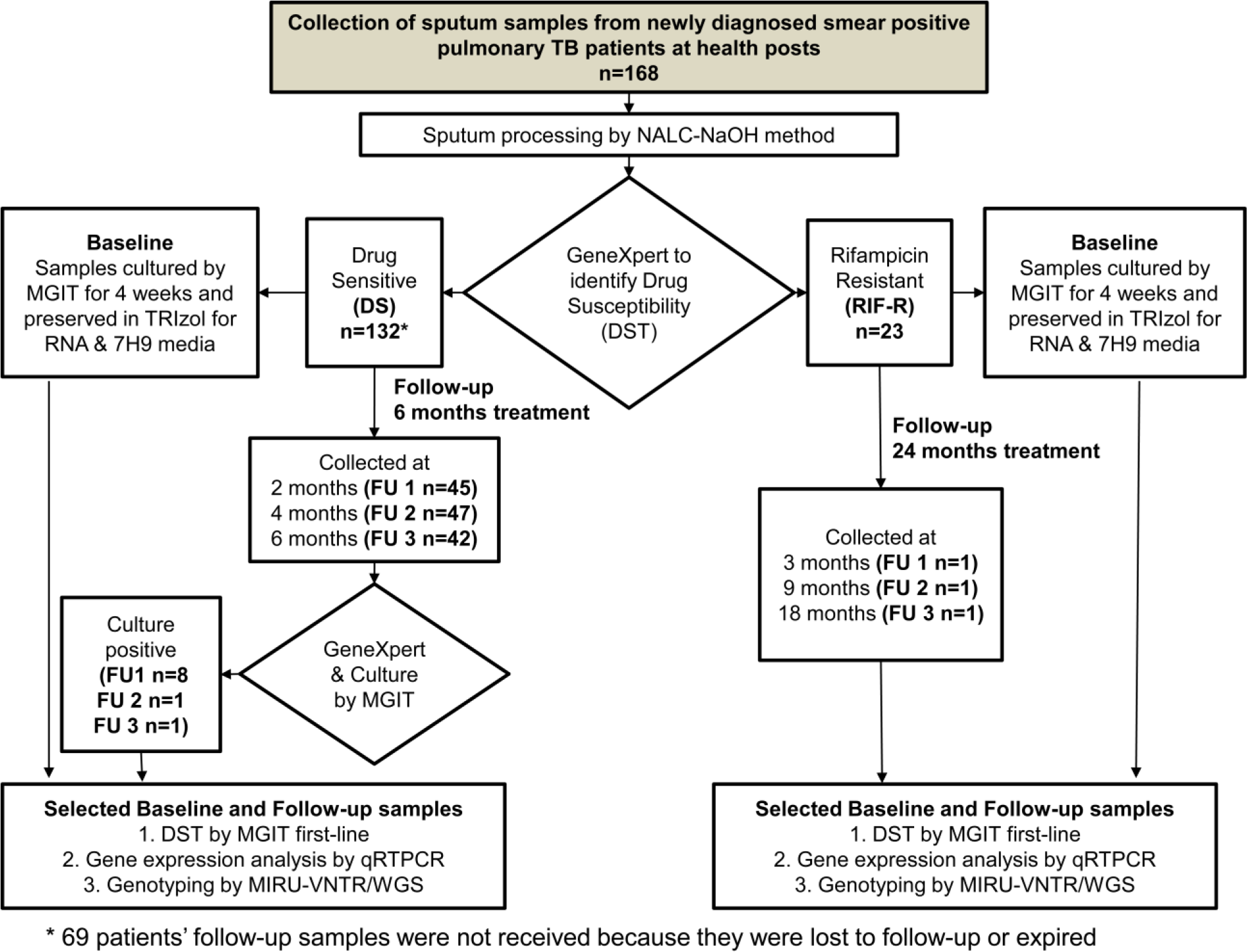
Schematic overview of experimental set-up

### Sample Processing and Drug susceptibility testing

The samples (baseline and follow-up) were processed using standard N-Acetyl-L-Cysteine (NALC) – Sodium Hydroxide (NALC-NaOH) method. The sputum concentrate was cultured by BACTEC™ MGIT™ 960 Mycobacteria Culture System (BD, Missouri, USA) as per manufacturer’s instructions. Cultures that were flagged positive in MGIT, were incubated up to 4 weeks and the resultant cultures were preserved in TriZol™ for total RNA extraction, as pellet for DNA isolation and in 20% glycerol for archiving. Phenotypic drug susceptibility testing (DST) was done for first-line anti-TB drugs Streptomycin, Isoniazid, Rifampicin and Ethambutol using BACTEC™ MGIT™-SIRE kit (BD, Missouri, USA) as per manufacturer’s instructions. The samples tested were clinical isolates that were culture positive at follow-up and their respective baseline pre-treatment isolates.

### Quantitative-Real Time Polymerase Chain Reaction (qRTPCR)

RNA extraction, cDNA preparation and qRTPCR were carried out as described earlier[10]. Primers for *ppsD, embC, Rv1457c, Rv1687c* and *recB* were same as described in the earlier study[10]. Each sample was tested in duplicate and the experiment was repeated three times from multiple cDNA preparations. The fold change was calculated with respect to standard laboratory strain H37Rv using ΔΔCt method and *fgd1* as the house-keeping gene.

### Statistical Analysis of qRTPCR data

Each group of data was subjected to normality test by the Shapiro Wilk test. The statistical significance for the study was calculated using Mann Whitney test. A p-value of less than 0.05 was considered significant. Both Mann Whitney and Shapiro Wilk test were performed using IBM-SPSS version 19 statistical software.

### Genotyping of clinical isolates

Genotyping was carried out to establish if the patient samples had the same strain of Mtb at baseline and follow-up. MIRU-VNTR (Mycobacterial interspersed repeat units-variable tandem repeats) was used for establishing the identity. For samples that gave inconclusive results by MIRU-VNTR, whole genome sequencing (WGS) was used to establish identity. MIRU-VNTR was carried out for 12-loci as described by Supply *et. al*[11] with minor modifications. The PCR for 12 loci was carried out individually using 12 sets of primers and conditions described by Supply *et. al.[11]* The resultant PCR fragments were analyzed by 1% agarose gel electrophoresis and the size estimated using gel analyzer with at least three lanes of 1Kb ladder as a marker. The estimated fragment size was assigned MIRU-VNTR alleles using Table-1 from Additional file 2 as described in Ali et. al [12]. The alleles were analyzed using MIRU-VNTR plus web-based platform for strain identity. The analysis was performed when data for all 12 loci were available. For whole genome sequencing, DNA was extracted from the preserved pellet as described in Chatterjee et al [13]. The sequencing was carried out by Genotypic Technologies Pvt Ltd, Bangalore on Illumina NextSeq 500 platform. The raw data in the form of FASTQ files were further used for strain identification using Mykrobe Predictor (v 0.4.2)^[14]^.

**Table-1.**
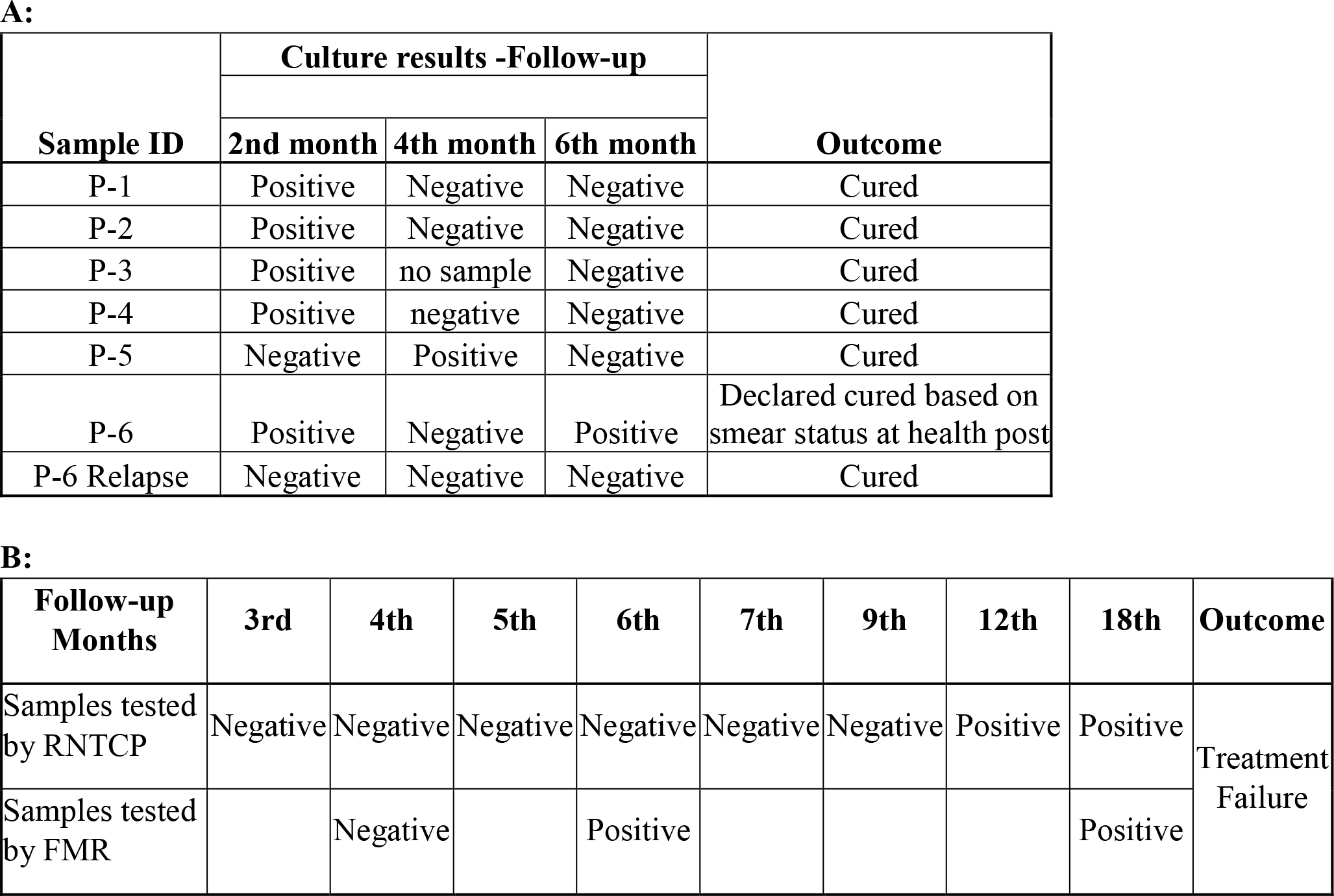
**A.** Culture results and outcome of baseline DS patients treated with 6 months first line anti-TB therapy. **B:** Culture result history of patient P7-Patient had MDR at baseline and on standardized 24 months CatIV treatment

### SDS Challenge Experiment

The experiment was carried out to investigate correlation between cell permeability of clinical strains with differing expression levels of *ppsD*. The permeability was measured as growth obtained in presence of a detergent sodium dodecyl sulfate (SDS) in the growth medium as compared to absence of detergent. *M.tb* clinical strains were grown to early log phase and a 10-fold dilution series with optical densities of 0.01 to 0.0001 at 580 nm was used. 5 μl of each dilution was spotted in triplicates onto 7H11 agar plates containing 10% oleic acid-albumin-dextrose-catalase with or without 0.01% SDS. The colonies were observed at 4 weeks after plating and the colony forming ability at different dilutions were compared in presence and absence of SDS.

## Results

Differential gene expression was studied for clinical isolates from patients with known treatment outcome information. The dropouts who could not be followed-up for outcome information were excluded. Of the 67 patients with known outcome information, 48 patients were cured (smear or culture negative at the end of treatment), 7 patients had completed treatment, 5 patients had delayed culture conversion but eventually cured (culture positive at first or second follow-up, but negative at the end of treatment) and 5 patients had poor outcome (treatment failure). Among the 5 poor outcome patients, 1 patient was culture positive, smear negative and drug sensitive at the end of treatment and had a relapse after four months which was cured; 4 patients acquired MDR during treatment as recorded by RNTCP. Two kinds of differential gene expression analysis were performed-First, patient samples were grouped based on treatment outcome/ resistance status and the differential gene expression analysis performed at baseline. Second, gene expression of culture positive follow-up samples that had identical strain as baseline was compared to the respective baseline expression and correlated to treatment response of the patient.

### *ppsD* expression is significantly associated with drug resistance

The patient samples were classified into four groups: **1.** Cured (n=18): Patients who were declared cured at the end of 6 months of treatment. All follow-up samples of these patients were culture negative or smear negative. **2.** Slow-responders (n=6): Patients who had at least one follow-up sample culture positive either at 2 or 4 months (delayed conversion) indicating a slower response to treatment, but eventually cured at the end of treatment (6 months). **3.** MDR converts (n=4): Four patients who were drug sensitive at baseline and acquired MDR during treatment. **4.** MDR at baseline (n=16): Patients who had MDR at baseline and were under Cat IV treatment for MDR-TB.

We compared gene expression levels of *ppsD, embC, recB, Rv1457c* and *Rv1687c* at baseline across these four groups. We observed that only *ppsD* showed differential expression in response to treatment. As evident from Fig 2, the median values of *ppsD* increased with respect to poor treatment response, *i.e* From 5.7 in the cured group to 8.3 in slow responders to 13 in MDR convert group. However, the change was statistically significant only between cured and MDR converts. Interestingly, the differential expression between the cured group and the patients who had MDR at baseline was also significantly different, suggesting an association of *ppsD* with the development of drug resistance. All other genes studied (*embC, recB, Rv1457c* and *Rv1687c)* did not show any significant effect with respect to treatment or drug resistance status (Fig 2). Only *Rv1457c* and *Rv1687c*, the efflux genes showed a trend similar to *ppsD*, between the groups, although not statistically significant. This suggests that alternate drug resistance mechanisms like cell permeability and efflux mechanism have an important role in patient’s response to treatment.

**Fig-2:**
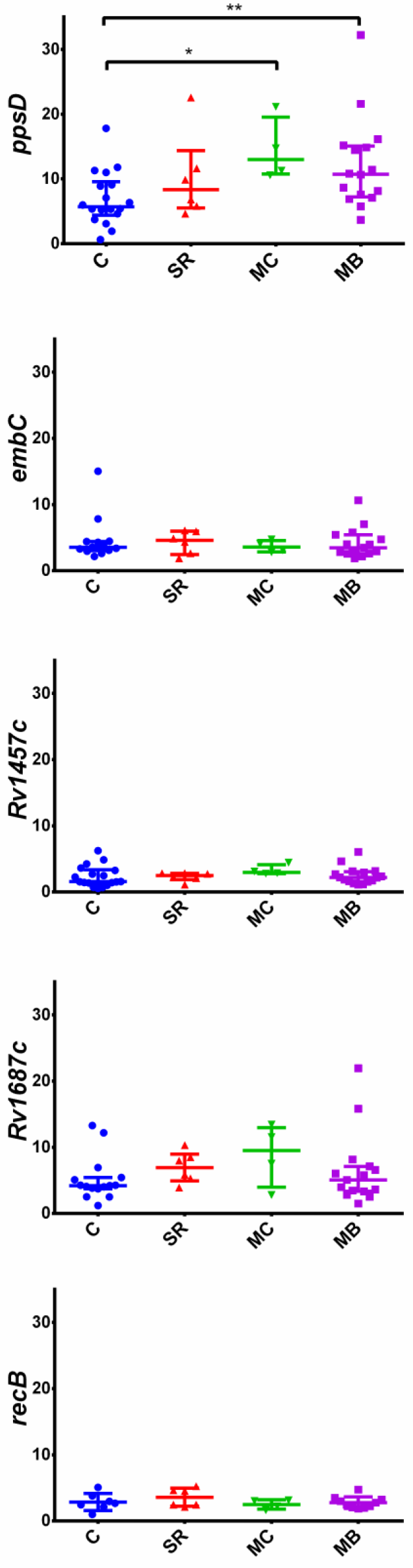
Comparison of gene expression in clinical isolates at baseline with treatment outcome and resistance status. Groups: 1. Cured [C] (n=18): Patients who were declared cured at the end of 6 months of treatment. All follow-up samples of these patients were culture negative and/or smear negative. 2. Slow-responders [SR] (n=6): Patients who had at least one follow-up sample culture positive indicating slower response to treatment, but eventually cured. 3. MDR converts [MC] (n=4): Four patients who were drug sensitive at baseline but acquired MDR during treatment. 4. MDR at baseline [MB] (n=16): Patients who had MDR at baseline and were under Cat IV treatment for MDR-TB.

### *ppsD* expression reflects treatment response in culture positive follow-up samples

To understand how gene expression varies with treatment, gene expression was tested in culture positive longitudinal isolates with same strain identity. The expression levels in the follow-up sample was compared to the baseline expression levels. We had three types of patients (Table 1A&B) who had varied culture positivity in follow-up resistance pattern and outcome-**1.**Patients (P1-5) who were drug sensitive and their initial follow-up samples were culture positive. Subsequent follow-ups were culture negative and defined cured (Table-1A). **2.**Patient (P6) who was drug sensitive all throughout, 1^st^ and 3^rd^ follow-up samples were culture positive although declared cured at 3^rd^ follow-up based on smear negative status at health post and ended the treatment at 6 months (Table-1A). This patient had a relapse of TB, 4 months after completion of the last treatment and eventually got cured during relapse treatment (all follow-ups during relapse were culture negative). **3.**Patient who had MDR at baseline (P7) and was on 24 months standardized MDR treatment. Although we did not get all follow-up samples of this patient, history of treatment response was tracked through data available from RNTCP and sporadic samples received by FMR (Table 1B). The patient’s final outcome was recorded as MDR treatment failure and the patient shifted to XDR treatment regimen.

As can be seen from Fig 3, in P1-5, *ppsD* expression was consistently down-regulated in follow-up samples compared to baseline in all patients (Fig -3). However, in the other genes studied, the change in expression levels were not consistent. -In both P6 and P7 patient samples, the expression of *ppsD, embC* and *Rv1457c* appeared to decrease with treatment initially (Fig 3). However, after a period of responding to treatment as seen by culture negative status (Table 1A-4^th^ month follow-up of P6, Table 1B-3^rd^ to 9^th^ month follow-up of P7), in the subsequent non-responsive state indicated by culture positivity (Table 1A-6^th^ month follow-up of P6, Table 1B-18^th^ month follow-up of P7), the expression levels of these 3 genes increased. Nevertheless, only *ppsD* showed treatment specific changes in all types of patients suggesting expression levels of *ppsD* reflects the patient’s response to treatment (Fig 3).

**Fig-3:**
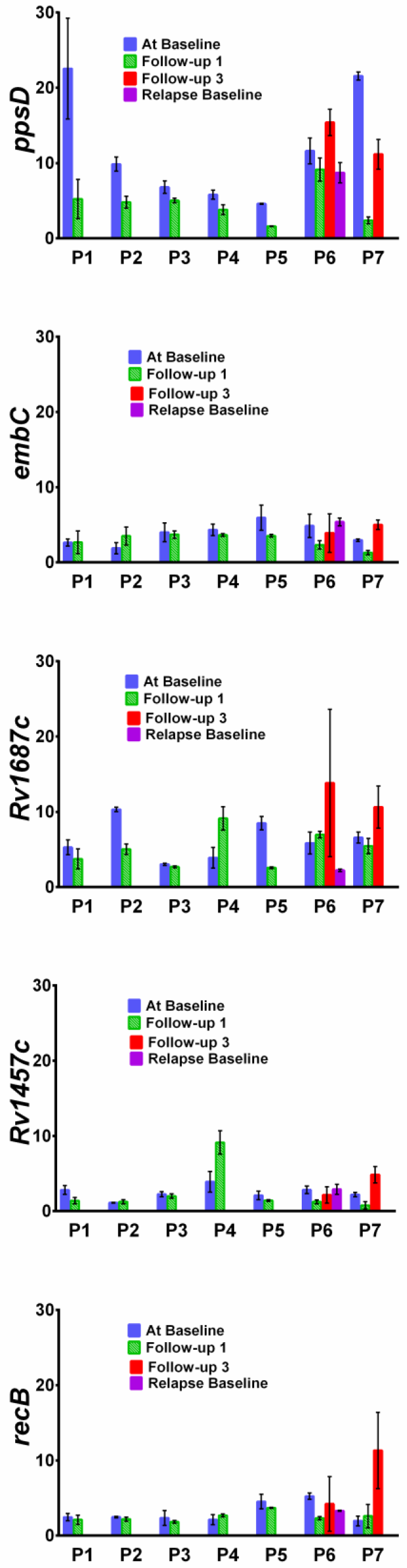
Comparison of *ppsD* expression in clinical isolates at baseline and follow-up with treatment response (culture result history). **P1-5** were DS patients whose initial follow-up samples were culture positive but subsequently were culture negative and cured. **P6** patient was drug sensitive throughout, 1st and 3rd follow-up samples were culture positive although declared cured at 3rd follow-up based on smear negative status at health post and ended the treatment at 6 months (Table-1A). The patient had a relapse of TB after 4 months. **P7** patient had MDR at baseline and was on standardized MDR treatment. the follow-up details are mentioned in Table-1B.

### Cell permeability and *ppsD* expression are not directly correlated

Since PDIM is critical for virulence, architecture and cell permeability[15,16], we hypothesized that *ppsD* expression levels may correlate to the level of cell permeability and thus influence treatment response. To check this, cell permeability of representative isolates from cured, slow responders, MDR converts and MDR at baseline groups were determined by their ability to grow in presence of detergent SDS. As evident from Fig 4, baseline isolates from different groups showed varying levels of growth retardation in presence of SDS (indicating higher permeable cell wall and hence more susceptibility to treatment). No direct correlation was observed between ability to grow on SDS with *ppsD* expression levels or treatment response. These results indicate that alterations in cell permeability alone may not explain the association of *ppsD* expression with treatment response in patients.

**Fig 4:**
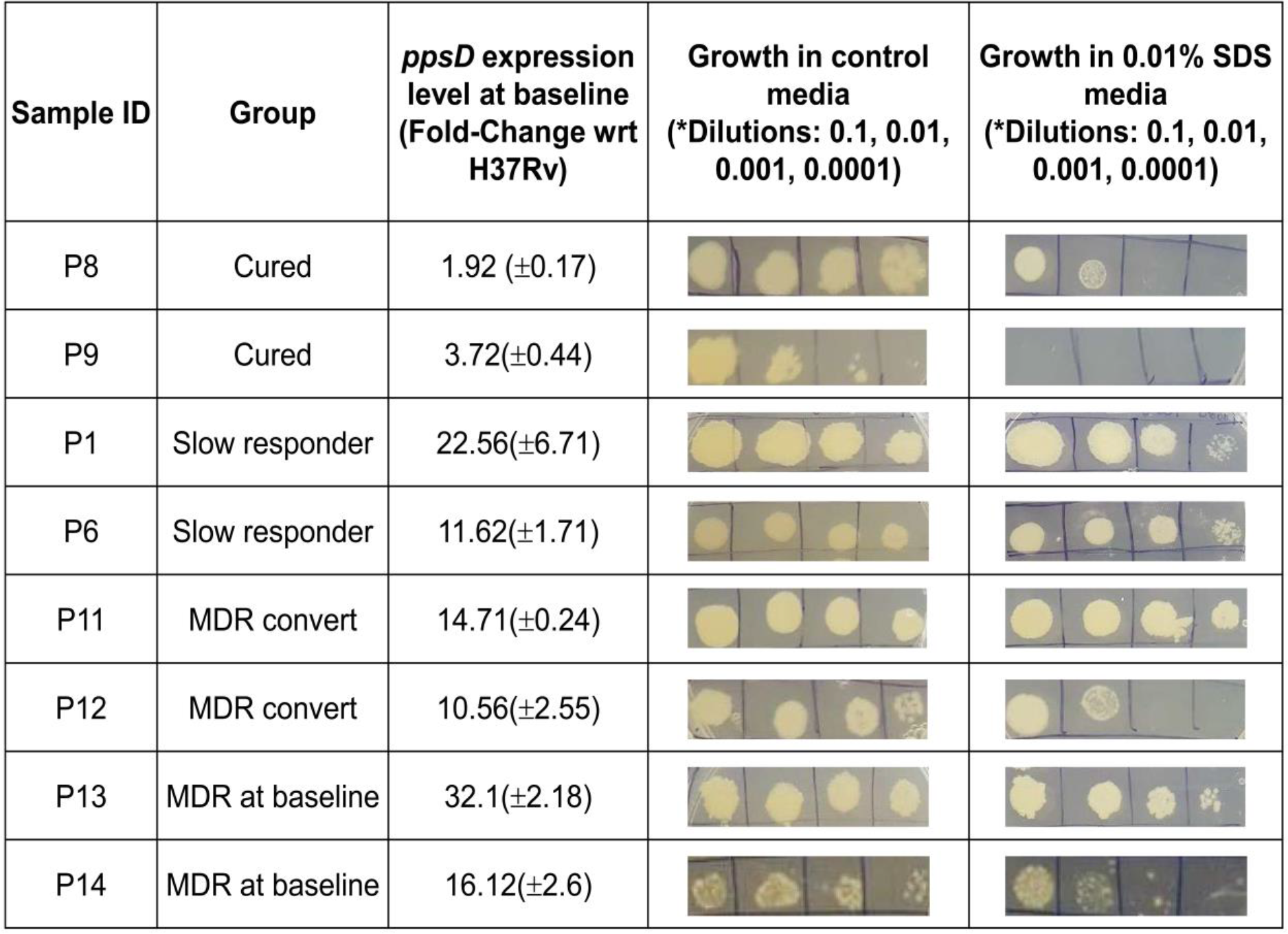
Growth of representative clinical isolates from different groups in presence of SDS detergent.

## Discussion

In the current study, we demonstrated that expression of *ppsD*, the gene involved in the synthesis of cell wall lipid PDIM, was significantly increased in patients with unfavourable treatment response. The isolates from patients who developed resistance during treatment (MDR convert) had higher *ppsD* expression than slow responders (delayed culture converters). Likewise, slow responders had higher *ppsD* expression than responsive cured patients (Fig 2). Even in isolates from patients who had MDR at baseline, the *ppsD* expression was significantly higher compared to cured patients. The higher expression of *ppsD* in MDR converts and baseline MDR group suggest that *ppsD* is associated with the development of drug resistance corroborating the observations from the earlier *in vitro* study[10]. In addition, the study shows that the *ppsD* expression levels correlate to treatment response in individual patients during treatment (Fig 3). The *ppsD* expression decreased in comparison to baseline values when culture positive isolates were collected prior to the period of culture negativity (responsiveness to treatment) in patients. However, *ppsD* expression levels increased to levels comparable to baseline in isolates of patients who showed a period of non-responsiveness (culture positivity) subsequent to a period of responsiveness to treatment (culture negativity) as seen in patients P6 and P7. This was observed irrespective of the resistance status of the isolates (P6 being DS and P7 being MDR). These results indicate that *ppsD* is involved in the development of drug resistance and response to treatment. Interestingly, higher expression of *ppsD* at baseline in patients with unfavourable outcome suggest that *ppsD* expression levels can potentially be used as a marker for predicting response to treatment in TB patients undergoing treatment irrespective of the drug resistance status.

*ppsD* gene is one of the five genes involved in the synthesis of PDIM, a long-chain-diol (phthiocerol) esterified with two branched-chain mycocerosic acids located in the outer mycobacterial cell wall that has been implicated in Mtb virulence[17]. PDIM has also been implicated in determining cell wall permeability in Mtb[16]. In our earlier study[10] it was suggested that differential expression of *ppsD* probably affected the levels of PDIM and hence the integrity of cell wall indicating propensity of strains with high *ppsD* expression to resist entry of drugs and allowing development of resistance through other mechanisms. Corroborating it, the DNA repair mechanisms and efflux pathways were are also observed to be deregulated. However in the current study, in patient isolates, although we observed expression levels of efflux genes following a pattern similar to *ppsD* expression at baseline (Fig 2 *Rv1687c* panel), the expression difference was not significant between the groups (cured, slow responders, MDR convert or MDR at baseline). In addition, the expression of efflux genes in longitudinal isolates did not show a “treatment response based” pattern similar to *ppsD*. Furthermore, SDS challenge experiments that determine cell permeability showed that there is no direct correlation between cell permeability and *ppsD* expression or treatment response. These results suggest that although the role of other mechanisms in treatment response and resistance development cannot be ruled out, *ppsD* may affect the treatment response of the Mtb to drugs through additional mechanisms. Studies have shown that strains that harbor mutation in *ppsD* leading to loss of PDIM synthesis are susceptible to interferon gamma (IFN-γ) mediated immunity [18]. It is possible that higher *ppsD* expression and hence higher levels of PDIM in strains resist IFN-γ mediated immunity in patients during treatment. Hence, a combination of both decreased permeability and increased resistance to the immune response of host may explain the differential treatment response in TB patients’ harbouring strains with differential *ppsD* expression.

To the best of our knowledge, this is the first study that demonstrates an association between differential gene expression and response to treatment in TB patients. Several studies have identified mutations in non-drug target genes that may serve as a compensatory mechanism for fitness of drug resistant strains[19,20]. Such mutations have been correlated to the mechanism of drug resistance but not to treatment response. The clinical isolates in our study may harbor such mutations that may explain differences in *ppsD* expression at baseline. Preliminary analysis of whole genome sequence carried out with isolates from our study did not yield any potential mutation in the promoter region or predicted regulators for *ppsD* expression (data not shown). However, the presence of indirect mutations that may control its expression could not be verified. In addition, in the current study, we have shown that the expression of *ppsD* varies with respect to treatment even in absence of observable drug resistance. Our results suggest that Mtb’s response to treatment is a complex phenomenon and may involve a combination of genetic changes and regulation of gene expression.

In conclusion, we have shown that *ppsD* expression varies with response to treatment in both DS and MDR patients and hence can potentially serve as a marker for predicting treatment response. However, the study was limited by the number of patient samples from whom complete sample and information were available due to high percentage of patients lost to follow-up. Hence, further research is warranted to confirm these findings in a larger cohort of patients and investigate its application as a predictor of treatment response. Moreover, *ppsD* is only one of the 8 genes that control PDIM synthesis. Hence this study raises several questions regarding the role of other genes involved in PDIM synthesis, how they correlate to treatment response and the genetic factors that determine the variability.

## Acknowledgements

This work was financially supported by Department of Science and Technology, Government of India [SB/SO/HS-0065/2013] and Navajbai Ratan Tata Trusts institutional grant to The Foundation for Medical Research (FMR). The authors thank Dr Ranjita Bagwe, Dr Thara Somashekar and their staff (Mumbai District TB Control Society, Mumbai) for collection of samples, Anil Dicholkar (FMR) for transport of samples, Shimoni Shah (FMR) for help with statistical analysis and Kayzad Nilgiriwala (FMR) for technical discussions.

## References

1. World Health Organization-Global Tuberculosis Report. http://apps.who.int/iris/bitstream/10665/259366/1/9789241565516-eng.pdf?ua=1 (2017).

2. TB India 2017-Revised National Tuberculosis Control Programme annual status report. https://tbcindia.gov.in/WriteReadData/TB%20India%202017.pdf (2017).

3. Dookie N, Rambaran S, Padayatchi N, Mahomed S, Naidoo K (2018) Evolution of drug resistance in Mycobacterium tuberculosis: a review on the molecular determinants of resistance and implications for personalized care. J Antimicrob Chemother 73 (5):1138–1151

4. Fonseca JD, Knight GM, McHugh TD (2015) The complex evolution of antibiotic resistance in Mycobacterium tuberculosis. Int J Infect Dis 32:94–100. doi:10.1016/j.ijid.2015.01.014

5. Louw GE, Warren RM, Gey van Pittius NC, McEvoy CRE, Van Helden PD, Victor TC (2009) A Balancing Act: Efflux/Influx in Mycobacterial Drug Resistance. Antimicrobial Agents and Chemotherapy 53 (8):3181–3189. doi:10.1128/aac.01577-08

6. Pule CM, Sampson SL, Warren RM, Black PA, van Helden PD, Victor TC, Louw GE (2015) Efflux pump inhibitors: targeting mycobacterial efflux systems to enhance TB therapy. J Antimicrob Chemother 71 (1):17–26

7. Machado D, Coelho TS, Perdigao J, Pereira C, Couto I, Portugal I, Maschmann RA, Ramos DF, von Groll A, Rossetti MLR, Silva PA, Viveiros M (2017) Interplay between Mutations and Efflux in Drug Resistant Clinical Isolates of Mycobacterium tuberculosis. Frontiers in microbiology 8:711. doi:10.3389/fmicb.2017.00711

8. Chatterjee A, Saranath D, Bhatter P, Mistry N (2013) Global transcriptional profiling of longitudinal clinical isolates of Mycobacterium tuberculosis exhibiting rapid accumulation of drug resistance. PloS one 8 (1):e54717. doi:10.1371/journal.pone.0054717

9. Penuelas-Urquides K, Gonzalez-Escalante L, Villarreal-Trevino L, Silva-Ramirez B, Gutierrez-Fuentes DJ, Mojica-Espinosa R, Rangel-Escareno C, Uribe-Figueroa L, Molina-Salinas GM, Davila-Velderrain J, Castorena-Torres F, Bermudez de Leon M, Said-Fernandez S (2013) Comparison of gene expression profiles between pansensitive and multidrug-resistant strains of Mycobacterium tuberculosis. Curr Microbiol 67 (3):362–371. doi:10.1007/s00284-013-0376-8

10. Sriraman K, Nilgiriwala K, Saranath D, Chatterjee A, Mistry N (2018) Deregulation of genes associated with alternate drug resistance mechanisms in Mycobacterium tuberculosis. Curr Microbiol 75 (4):394–400

11. Supply P, Lesjean S, Savine E, Kremer K, Van Soolingen D, Locht C (2001) Automated high-throughput genotyping for study of global epidemiology of Mycobacterium tuberculosis based on mycobacterial interspersed repetitive units. J Clin Microbiol 39 (10):3563–3571

12. Ali A, Hasan Z, Tanveer M, Siddiqui AR, Ghebremichael S, Kallenius G, Hasan R (2007) Characterization of Mycobacterium tuberculosis Central Asian Strain1 using mycobacterial interspersed repetitive unit genotyping. BMC Microbiol 7 (1):76

13. Chatterjee A, Nilgiriwala K, Saranath D, Rodrigues C, Mistry N (2017) Whole genome sequencing of clinical strains of Mycobacterium tuberculosis from Mumbai, India: A potential tool for determining drug-resistance and strain lineage. Tuberculosis 107:63–72

14. Bradley P, Gordon NC, Walker TM, Dunn L, Heys S, Huang B, Earle S, Pankhurst LJ, Anson L, De Cesare M (2015) Rapid antibiotic-resistance predictions from genome sequence data for Staphylococcus aureus and Mycobacterium tuberculosis. Nature communications 6:10063

15. Goude R, Parish T (2008) The genetics of cell wall biosynthesis in Mycobacterium tuberculosis. Future Microbiology 3:299–313

16. Camacho LR, Constant P, Raynaud C, Lanéelle M-A, Triccas JA, Gicquel B, Daffé M, Guilhot C (2001) Analysis of the Phthiocerol Dimycocerosate Locus ofMycobacterium tuberculosis: Evidence that this lipid is involved in the cell wall permeability barrier. J Biol Chem 276 (23):19845–19854

17. Camacho LR, Ensergueix D, Perez E, Gicquel B, Guilhot C (1999) Identification of a virulence gene cluster of Mycobacterium tuberculosis by signature‐tagged transposon mutagenesis. Mol Microbiol 34 (2):257–267

18. Kirksey MA, Tischler AD, Siméone R, Hisert KB, Uplekar S, Guilhot C, McKinney JD (2011) Spontaneous phthiocerol dimycocerosate-deficient variants of Mycobacterium tuberculosis are susceptible to gamma interferon-mediated immunity. Infect Immun 79 (7):2829–2838

19. Al-Saeedi M, Al-Hajoj S (2017) Diversity and evolution of drug resistance mechanisms in Mycobacterium tuberculosis. Infection and drug resistance 10:333

20. Farhat MR, Shapiro BJ, Kieser KJ, Sultana R, Jacobson KR, Victor TC, Warren RM, Streicher EM, Calver A, Sloutsky A (2013) Genomic analysis identifies targets of convergent positive selection in drug-resistant Mycobacterium tuberculosis. Nat Genet 45 (10):1183–1189

